# Characterizing Resting-State Brain Dynamics with Frequency-Resolved EEG Microstates: Parallel Analyses of Psilocybin Microdosing and Acute Inhaled DMT

**DOI:** 10.64898/2026.05.05.723034

**Authors:** Povilas Tarailis, Inga Griskova-Bulanova

**Affiliations:** eBrain Lab, School of Mechatronic Systems Engineering, Simon Fraser University, Surrey, British Columbia, Canada; Faculty of Medicine, Translational Health Research Institute, Vilnius University, Vilnius, Lithuania

## Abstract

Electroencephalographic (EEG) microstates provide a compact framework for characterizing the temporal organization of large-scale brain activity, yet their sensitivity to altered brain states remains insufficiently explored. In this study, we applied broadband and frequency-resolved EEG microstate analysis to resting-state EEG data from two publicly available datasets acquired under markedly different altered-state conditions: psilocybin microdosing and acute inhaled N,N-dimethyltryptamine (DMT). The aim was to determine whether narrowband microstate analysis reveals structured alterations in resting-state brain dynamics beyond those captured by broadband analysis alone. Psilocybin microdosing was associated with relatively subtle effects, including reduced global field power and frequency-specific alterations in delta- and theta-band microstate parameters, while no significant broadband spatiotemporal changes were observed. In contrast, acute inhaled DMT was associated with broader microstate alterations spanning broadband, delta, theta, and alpha activity, indicating more extensive reorganization of temporal microstate expression. Across both datasets, a descriptive overlap was observed in the delta band, where microstate C showed increased duration and microstate D showed decreased occurrence. Given the substantial differences between datasets in dose, route of administration, temporal dynamics, and study context, these overlapping effects should be interpreted cautiously. Overall, the findings support frequency-resolved EEG microstate analysis as a useful approach for characterizing altered resting-state brain dynamics and for detecting frequency-specific effects that may be obscured in broadband summaries.

## Introduction

Electroencephalographic (EEG) microstates are brief periods of quasi-stable scalp potential topographies that are thought to reflect transient large-scale functional states of the brain (Michel and Koenig, 2018; Tarailis et al., 2024a). Rather than describing brain activity solely in terms of oscillatory power, microstate analysis captures the temporal segmentation of ongoing EEG into recurring spatial configurations and thus provides a compact framework for studying the spatiotemporal organization of whole-brain dynamics.

Microstate analysis has become an increasingly important tool for investigating large-scale brain organization in both healthy and clinical populations (Haydock et al., 2025; Michel and Koenig, 2018; Tarailis et al., 2024a). Resting-state microstate dynamics have been linked to fMRI-derived functional networks (Britz et al., 2010; Custo et al., 2017), and alterations in microstate parameters have been associated with variation across cognitive, affective, and clinical domains (Tarailis et al., 2024a). However, conventional microstate analysis is typically performed on broadband EEG, which may obscure frequency-specific aspects of brain-state organization.

Recent studies suggest that this limitation can be partly addressed by narrowband microstate analysis, in which EEG is decomposed into conventional frequency ranges prior to segmentation (Férat et al., 2022; Terpou et al., 2022). Although narrowband and broadband approaches tend to yield broadly similar topographic classes, their temporal characteristics may differ substantially, indicating that band-limited analysis can reveal aspects of brain dynamics that remain hidden in broadband summaries (Férat et al., 2022; Terpou et al., 2022; Xue et al., 2024). This is particularly relevant when expected effects are subtle, frequency-dependent, or heterogeneously expressed across large-scale networks. A combined analysis of broadband and narrowband microstates may therefore provide a more informative description of altered resting-state brain dynamics than spectral power estimates alone.

Psychedelic states offer a useful context in which to examine the sensitivity of such measures. Classical psychedelics such as psilocybin and N,N-dimethyltryptamine (DMT) produce marked alterations in conscious experience and have become an important model for studying altered brain states (Nichols et al., 2017). At the pharmacological level, these compounds share prominent serotonergic actions, particularly involving 5-HT_2_A receptors, but differ substantially in dose, route of administration, temporal dynamics, and experiential intensity (Hatzipantelis and Olson, 2024; Nichols et al., 2017). EEG studies have most consistently reported changes in oscillatory activity in the alpha range, whereas findings in other frequency bands have been more heterogeneous across studies and experimental settings (Kometer et al., 2013; Skosnik et al., 2023; Timmermann et al., 2019). Much less is known about how these conditions affect the temporal organization of large-scale EEG states as indexed by microstate dynamics.

In the present study, we applied broadband and frequency-resolved EEG microstate analysis to two publicly available resting-state datasets: one collected after psilocybin microdosing (Cavanna et al., 2022) and one collected during acute inhaled DMT effects (Tagliazucchi et al., 2021). These datasets differ markedly in dose, route of administration, temporal profile, and study context (Good et al., 2023; Madsen et al., 2019), and therefore do not support a direct substance comparison. Instead, they provide two contrasting perturbation regimes allowing to test whether frequency-resolved microstate analysis is sensitive to structured alterations in resting-state brain dynamics. Examining both datasets in parallel therefore allows assessment of whether broadband and narrowband microstate measures capture overlapping or distinct patterns across substantially different psychedelic conditions. The aim of this study was therefore to determine 1) whether frequency-resolved microstate analysis detects band-specific alterations in resting-state brain-state dynamics across two independent psychedelic datasets, 2) whether narrowband decomposition provides information beyond broadband microstate analysis, and 3) whether the observed effects reflect shared or dataset-specific patterns.

## Methods and materials

The study workflow is summarised in Figure 1.

**Figure 1.**
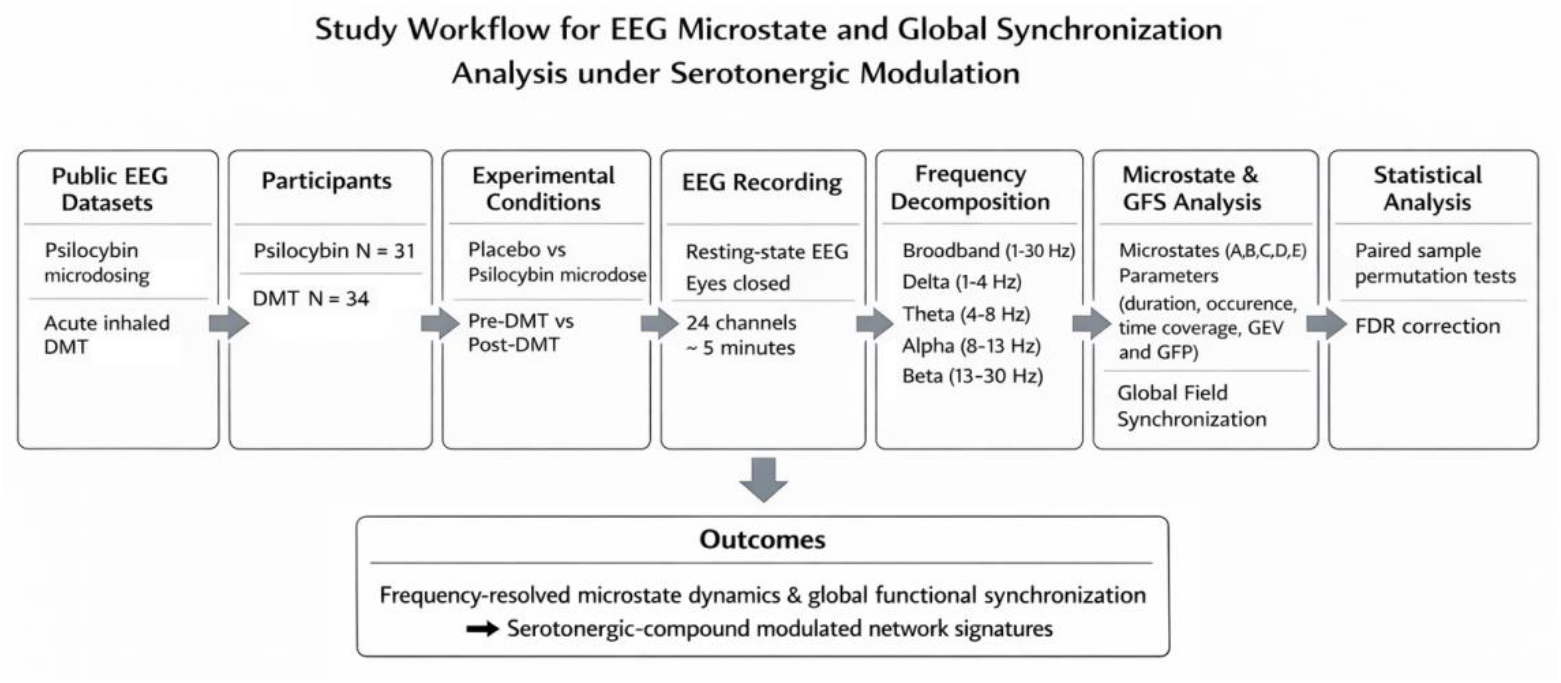
A schematic representation of the study workflow.

### Study design and datasets

This study is a secondary analysis of two previously published, publicly available resting-state EEG datasets acquired under psychedelic conditions. One dataset examined psilocybin microdosing and the other acute inhaled DMT. Raw EEG files were downloaded from public repositories (psilocybin: OSF, https://osf.io/hnxq6/; DMT: Zenodo, https://zenodo.org/records/3992359). Full details of recruitment, inclusion criteria, and original study procedures are reported in Cavanna et al. (2022) and Tagliazucchi et al. (2021). Both original studies were conducted in accordance with the Declaration of Helsinki, approved by local research ethics committees, and based on written informed consent from all participants.

### Participants

The psilocybin dataset included 34 participants (11 females, 23 males; mean age 31.26 ± 4.41 years). On average, participants reported 11 ± 14.9 previous psychedelic experiences. The DMT dataset included 35 participants (7 females, 28 males; mean age 33.1 ± 6 years), with an average of 92.2 ± 201.4 experiences with ayahuasca and 3.6 ± 5.6 experiences with DMT alone.

### Drug administration

In the psilocybin dataset, participants took part in a randomized, double-blind, placebo-controlled crossover procedure within the broader observational study framework. Each participant completed two study weeks, receiving an active capsule during one week and a placebo capsule during the other. The active capsule contained 0.5 g of ground dried psilocybin mushrooms, while the placebo capsule contained edible mushroom material. Active and placebo conditions were administered in randomized order, and both participants and researchers were blind to capsule identity. According to alkaloid analysis reported in the original study, the mushroom material contained 640.2 μg/g psilocybin and 950.7 μg/g psilocin, corresponding to an estimated total of approximately 0.8 mg psilocybin + psilocin per 0.5 g dose (Cavanna et al., 2022).

In the DMT dataset, participants inhaled freebase DMT in a naturalistic setting using a pipe. The pipe was withdrawn when participants leaned back or when its contents were exhausted. Each session involved an estimated dose of 40 mg freebase DMT, as reported in the original study.

### Subjective data

Subjective data were obtained from the original studies. In the psilocybin dataset, subjective drug effects were assessed using a visual analog scale (VAS) based on 21 items adapted from Carhart-Harris et al. (2016), and total scores were used in the present analysis. In the DMT dataset, participants completed several post-experience questionnaires, including the 5D altered states of consciousness scale (5D-ASC), the Mystical Experience Questionnaire (MEQ30), and the Near-Death Experience scale. The present analysis used total 5D-ASC scores only.

### EEG recordings

EEG signals were acquired using 24 Ag/AgCl electrodes mounted in an elastic cap (EASYCAP GmbH, Inning, Germany) and recorded with a portable research-grade mobile EEG system (mBrainTrain LLC, Belgrade, Serbia) attached to the back of the cap between O1 and O2. Reference and ground electrodes were placed at FCz and AFz, respectively. Data were sampled at 500 Hz using an online 0– 250 Hz passband filter.

In the psilocybin dataset, five minutes of eyes-closed EEG were recorded approximately 1.5 hours after drug administration. In the DMT dataset, eyes-closed EEG was recorded for 5 minutes before DMT administration and again immediately after participants exhaled DMT vapor, continuing until they indicated a return to baseline (mean duration 6 ± 1.4 min).

### EEG data preprocessing

Both datasets were preprocessed using the same pipeline in MATLAB and the EEGLAB toolbox (Delorme and Makeig, 2004). EEG data were filtered using a two-pass, zero-phase finite impulse response bandpass filter (1–30 Hz). Non-cerebral artifacts, including muscle activity, eye movements, cardiac pulse, and jaw tension, were corrected using independent component analysis (ICA). Channels with excessive artifacts were rejected and reconstructed using spherical spline interpolation, and data were then re-referenced to the average reference.

Records from three participants in the psilocybin dataset and one participant in the DMT dataset were excluded because of missing or corrupted files, resulting in final samples of 31 and 34 participants, respectively.

For each participant and condition, EEG data were retained as broadband signals (1–30 Hz) and additionally filtered into delta (1–4 Hz), theta (4–8 Hz), alpha (8–13 Hz), and beta (13–30 Hz) frequency ranges, following previous narrowband microstate studies (Férat et al., 2022; Terpou et al., 2022; Xue et al., 2024). This yielded one broadband and four narrowband datasets for subsequent microstate analysis.

### Microstate segmentation and backfitting

Microstate segmentation was performed using Cartool (v5.01) (Brunet et al., 2011) at both the individual and group levels for each dataset and frequency range. For each participant, scalp topographies at peaks of global field power (GFP) were extracted and submitted to k-means clustering to identify dominant individual topographies. The clustering procedure was repeated 50 times while increasing the number of possible cluster solutions from 1 to 10. The optimal number of clusters for each participant was determined using the meta-criteria implemented in Cartool (Artoni and Michel, 2025; Brunet et al., 2011).

At the group level, the dominant individual topographies were concatenated across participants and submitted to a second k-means clustering procedure. The optimal number of group-level microstate classes was again selected using the Cartool meta-criteria. Group-level templates were derived jointly across conditions within each dataset so that between-condition comparisons were performed on a common set of topographies. This approach emphasizes condition-related differences in the temporal expression of shared maps rather than differences arising from separately estimated condition-specific templates (Murphy et al., 2024).

The resulting group-level templates were backfitted to individual EEG data using a winner-takes-all procedure based on spatial correlation. Polarity was ignored, and a minimum correlation threshold was applied. Temporal smoothing was performed using a 20 ms window with a smoothing factor of 10 to reduce spurious short segments during low-GFP periods. Segments lasting 20 ms or less were reassigned to neighboring segments.

For each microstate class, condition, and frequency range, the following parameters were extracted: duration, defined as the mean uninterrupted segment length; occurrence, defined as the number of segments per second; time coverage, defined as the percentage of total analysis time occupied by a given microstate; global explained variance (GEV), defined as the proportion of EEG topographic variance explained by a given microstate class weighted by GFP; and mean global field power (GFP), reflecting the average field strength associated with that microstate class.

### Statistical analysis

Microstate parameters were compared between conditions separately for each dataset and frequency range using paired-sample permutation tests (Proschan et al., 2014). In the psilocybin dataset, comparisons were made between active and placebo conditions. In the DMT dataset, comparisons were made between pre-DMT and post-DMT conditions. P values were estimated using 10,000 random permutations.

To control for multiple comparisons, false discovery rate (FDR) correction (Benjamini and Hochberg, 1995) was applied within each dataset and frequency range across the full set of tested microstate-parameter comparisons. A corrected significance threshold of α = 0.05 was used. Effect sizes are reported as Hedges’ g (Durlak, 2009; Lakens, 2013).

Associations between subjective ratings and microstate parameters were assessed using Pearson correlation coefficients. In the psilocybin dataset, correlations were computed with total VAS scores; in the DMT dataset, correlations were computed with total 5D-ASC scores. Correlation p values were corrected using the FDR procedure.

## Results

### Microstate topographies

Group-level microstate topographies for both datasets are shown in Figure 2B. In the psilocybin dataset, group templates were derived from the combined placebo and psilocybin conditions within each frequency range. The optimal number of microstate classes was five across all frequency ranges, and the resulting topographies corresponded to prototypical spatial configurations consistent with previously described meta-microstates and commonly reported microstate classes (Koenig et al., 2024; Tarailis et al., 2024a). In the DMT dataset, group templates were derived from the combined pre- and post-DMT conditions within each frequency range. The optimal number of classes was also five in the broadband, delta, and theta ranges, whereas a sixth microstate was identified in the alpha and beta ranges. This additional microstate, characterized by a symmetric left-to-right configuration, is consistent with patterns reported in previous EEG microstate studies (Bréchet et al., 2019; Custo et al., 2017; Koenig et al., 2024).

**Figure 2.**
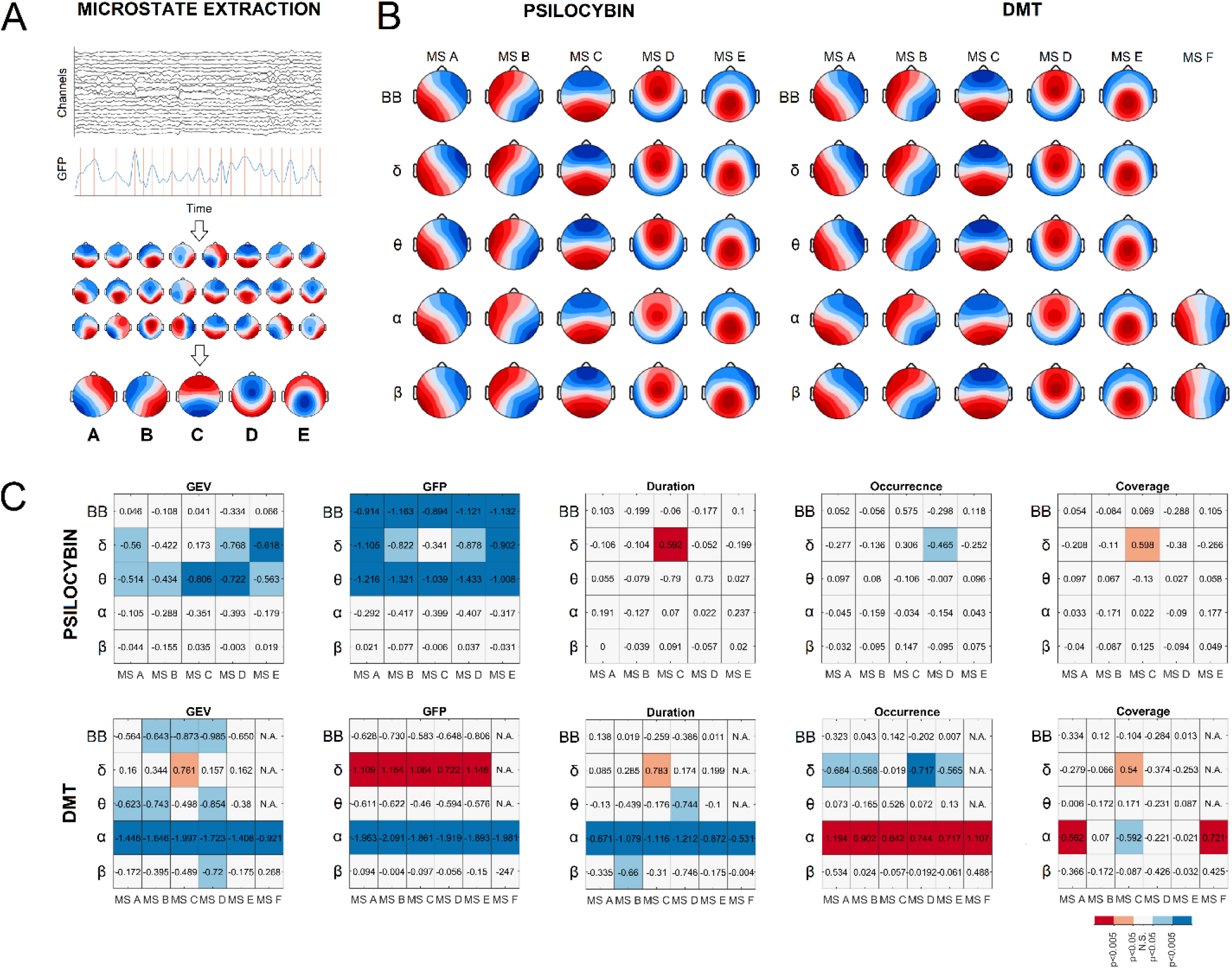
A.A schematic representation of the microstate approach and microstate A, B, C, D, E class topographies. B. EEG microstate topographies obtained at different frequency ranges (BB - broadband, δ, θ, α, and β) in the Psilocybin and DMT datasets. C. Hedges’ g effect sizes for comparisons between Psilocybin and Placebo, and Pre- and Post-DMT conditions across extracted microstate parameters. Each box represents the effect size for a given parameter. Red boxes indicate a significant increase following drug administration (FDR corrected p < 0.05), blue boxes indicate a significant decrease, and white boxes denote non-significant differences between conditions.

### Microstate parameters

#### Psilocybin dataset

The effect sizes for condition comparisons in the psilocybin dataset are shown in Figure 2C. Descriptive statistics (means and standard deviations) for the extracted parameters under psilocybin and placebo conditions are provided in Supplementary Table S1. No significant differences in broadband spatiotemporal microstate parameters, including GEV, duration, occurrence, and time coverage, were observed between placebo and psilocybin conditions. However, GFP was significantly reduced after psilocybin administration across all microstate classes (p < 0.005, Hedges’ g < -0.894). Narrowband analysis revealed similar GFP reductions in the theta (p < 0.05, Hedges’ g < -1.008) and delta (p < 0.05, Hedges’ g < -0.822) frequency ranges, except for microstate D in the delta band, for which no significant GFP change was observed. Additional narrowband microstate-specific differences were also observed between placebo and psilocybin conditions. Following psilocybin administration, GEV was reduced for microstates A, D, and E in the delta range (p < 0.05, Hedges’ g < -0.560), as well as across all microstate classes in the theta range (p < 0.05, Hedges’ g < -0.514). In the delta band, microstate D showed significantly decreased occurrence (p < 0.05, Hedges’ g = - 0.465), whereas microstate C showed increased mean duration (p < 0.005, Hedges’ g = 0.592) and increased time coverage (p < 0.05, Hedges’ g = 0.598). No significant differences in microstate parameters were observed in the alpha or beta frequency ranges. After correction for multiple comparisons, no significant correlations were found between microstate parameters and individual VAS scores across any frequency range (Supplementary Table S3).

### DMT dataset

The effect sizes for condition comparisons in the DMT dataset are shown in Figure 2C. Descriptive statistics (means and standard deviations) for the extracted parameters under pre- and post-DMT conditions are provided in Supplementary Table S2. Relative to the psilocybin dataset, the DMT dataset showed broader alterations across temporal and power-related microstate parameters. In the broadband EEG, significant reductions in GEV were observed for microstates B, C, and D following DMT administration (p < 0.05, Hedges’ g < -0.643). DMT also induced specific alterations in delta-range microstate dynamics. In this frequency band, microstate C showed increased GEV (p < 0.05, Hedges’ g = 0.761), increased mean duration (p < 0.05, Hedges’ g = 0.783), and increased time coverage (p < 0.05, Hedges’ g = 0.540). In contrast, the remaining four microstates showed reduced occurrence rates (p < 0.05, Hedges’ g < -0.565). Additionally, GFP was elevated across all microstates in the delta range following DMT administration compared with the pre-DMT condition (p < 0.005, Hedges’ g > 0.722). In the theta range, reductions in GEV were observed for microstates A, B, and D (p < 0.05, Hedges’ g < -0.623), together with a significant decrease in the duration of microstate D (p < 0.05, Hedges’ g = -0.744). In the alpha range, all microstates showed decreased GEV (p < 0.005, Hedges’ g < -0.921), decreased GFP (p < 0.005, Hedges’ g < -1.861), and decreased mean duration (p < 0.005, Hedges’ g < -0.531), alongside increased occurrence rates (p < 0.005, Hedges’ g < 1.194). In addition, time coverage increased for microstates A and E (p < 0.05, Hedges’ g = 0.562 and p < 0.05, Hedges’ g = 0.721, respectively), while it decreased for microstate C (p < 0.05, Hedges’ g = -0.592). For beta range, a decrease in GEV of MS D (p < 0.05, Hedges’ g = -0.720) and duration of MS B (p < 0.05, Hedges’ g = -0.660) were observed. After correction for multiple comparisons, no significant correlations were found between microstate parameters and 5D-ASC scores (Supplementary Table S4).

## Discussion

The present study examined whether frequency-resolved EEG microstate analysis can detect structured alterations in resting-state brain dynamics beyond those captured by broadband segmentation alone. Across two independent datasets, narrowband microstate analysis was sensitive to condition-related changes, but the extent and distribution of these changes differed substantially. In the psilocybin microdosing dataset, effects were relatively subtle and concentrated in the delta and theta ranges. In the acute inhaled DMT dataset, alterations were broader and extended across broadband, delta, theta, and alpha activity. These findings support the utility of narrowband microstate decomposition for characterizing altered resting-state brain dynamics, particularly when effects are frequency-specific or unevenly distributed across parameters (Férat et al., 2022; Terpou et al., 2022; Xue et al., 2024).

### Narrowband microstates reveal effects beyond broadband analysis

A central methodological observation is that broadband and narrowband analyses were not equally informative. In the psilocybin dataset, broadband spatiotemporal microstate parameters remained largely unchanged, despite a consistent reduction in broadband GFP across all microstate classes. Additional effects emerged only after decomposition into conventional frequency bands, with changes concentrated in the delta and theta ranges. This indicates that condition-related differences may remain largely undetected at the broadband spatiotemporal level while becoming visible in band-limited microstate parameters. Such a pattern is consistent with previous work showing that broadband and narrowband microstate segmentations may share similar topographic classes while differing in temporal expression (Férat et al., 2022; Terpou et al., 2022). The present results therefore reinforce the view that narrowband microstate analysis can increase sensitivity when effects are subtle or restricted to specific frequency ranges.

The DMT dataset extends this methodological point in a different way. Here, condition-related changes were already evident at the broadband level, but narrowband analysis provided a richer description of how those alterations were distributed across frequency ranges and microstate parameters. In particular, alpha-band microstates showed shorter mean duration, reduced explained variance, reduced field strength, and increased occurrence across all classes, indicating shorter-lived and more frequently recurring alpha-band microstates. Delta- and theta-range changes added further evidence that the altered state was not expressed uniformly across the spectrum. Thus, whereas the psilocybin dataset illustrates the increased sensitivity of narrowband decomposition to subtle effects, the DMT dataset shows how frequency-resolved microstate analysis can resolve broader reorganization into parameter- and frequency-specific patterns rather than a single undifferentiated broadband effect.

### Relation to prior findings from the same datasets

Narrowband microstate analysis revealed frequency-specific alterations that were broadly compatible with prior findings from the same datasets, while adding a complementary account of how these effects were expressed in the temporal organization of resting-state EEG.

In the psilocybin dataset, the original report described noticeable but limited acute subjective effects, reduced theta power, and preserved broadband signal complexity as assessed by Lempel-Ziv complexity (Cavanna et al., 2022). Against that background, the present microstate findings suggest that even when broadband complexity remains unchanged and spectral effects are limited, subtle condition-related differences may still be expressed at the level of temporal microstate organization, particularly in lower-frequency ranges. The added value of the present reanalysis is therefore not the claim of a qualitatively different psilocybin effect, but the demonstration that temporal microstate measures can reveal structured narrowband alterations in a low-intensity condition.

In the DMT dataset, prior analyses reported prominent alpha suppression together with changes in lower-frequency activity (Tagliazucchi et al., 2021; Timmermann et al., 2019, 2023). The present findings complement those results by shifting the focus from oscillatory power to the temporal segmentation of resting-state activity. Rather than simply confirming altered alpha or low-frequency power, the microstate analysis shows how the DMT condition is accompanied by changes in the stability, occurrence, and relative contribution of recurring scalp topographies over time. This complementary level of description may be especially useful when spectral and spatiotemporal measures do not move in parallel, as suggested here by altered theta microstate parameters alongside previously reported theta-power increases in the same dataset (Tagliazucchi et al., 2021).

### Patterns across contrasting perturbation regimes

Both datasets involve classical psychedelics with overlapping primary serotonergic targets; however, their substantial differences in dose, route of administration, and temporal profile mean that the present design is better suited to identifying convergent and divergent microstate patterns than to inferring a common mechanism. Although the two datasets should not be treated as a direct substance comparison, they are informative as contrasting perturbation regimes. The psilocybin dataset represents a low-intensity condition in which effects were largely confined to lower-frequency microstate measures, whereas the DMT dataset reflects a broader redistribution across several parameters and frequency bands. From a methodological perspective, this contrast is useful because it shows that frequency-resolved microstate analysis remains informative across markedly different effect magnitudes. In one case, it reveals changes that are largely absent from broadband spatiotemporal measures (Figure 2C); in the other, it resolves a broader pattern of reorganization into distinct frequency-specific components (Figure 2C).

A descriptive overlap was also observed in the delta band, where both datasets showed increased duration of microstate C and decreased occurrence of microstate D. This convergence is potentially relevant, but it should be interpreted cautiously. Given the substantial differences between datasets, the present design does not allow these overlapping effects to be attributed to compound identity or to a single shared mechanism. At most, the findings suggest that some low-frequency microstate features may recur across distinct perturbation regimes and may therefore warrant targeted evaluation in future work. Interpretation of these commonalities at the level of prototypical microstate function should likewise remain restrained. MS C has been associated in previous literature with altered patterns of external engagement and large-scale functional coordination in a range of conditions, including sleep, hypnosis, and psychosis-related states (Brodbeck et al., 2012; Katayama et al., 2007; Liebrand et al., 2024; Rieger et al., 2016). While this makes the repeated involvement of microstate C noteworthy, the present data are not sufficient to support strong inferences about shared experiential or neurophysiological mechanisms across the two datasets. The main implication is therefore methodological: frequency-resolved microstate analysis may detect recurring low-frequency features across different perturbation regimes, but their interpretation requires additional evidence.

### Subjective measures

No significant correlations were observed between microstate parameters and available subjective ratings in either dataset after correction for multiple comparisons. This absence of association should be interpreted cautiously. Subjective measures were limited to post hoc summary ratings and were therefore only loosely matched to the temporal scale of the EEG-derived measures. In addition, the available questionnaires differed across datasets, and the present analysis relied on total scores rather than finer-grained experiential dimensions. Under these conditions, weak or absent associations are not unexpected. Nevertheless, this result also differs from earlier work linking microstate measures to self-reported spontaneous thought and mind-wandering (Pipinis et al., 2016; Tarailis et al., 2021), as well as from prior analyses of the DMT dataset reporting associations between selected EEG measures and subjective reports (Pallavicini et al., 2021; Tagliazucchi et al., 2021). In the psilocybin parent study, the main reported neural findings were reduced theta power and preserved broadband signal complexity, accompanied by modest acute subjective effects that appeared strongly influenced by unblinding/expectancy rather than being directly associated with the EEG measures (Cavanna et al., 2022). Accordingly, the present null findings should not be treated as evidence against the relevance of microstate measures, but rather as an indication that the available subjective indices were not optimally matched to the temporal and descriptive scale of the EEG analysis. At present, the microstate alterations described here are best understood as changes in resting-state temporal organization rather than as validated electrophysiological proxies of subjective intensity.

## Limitations

Several limitations should be considered when interpreting the present findings. First, this was a secondary analysis of two previously published datasets that differed substantially in design, dose, route of administration, temporal dynamics, and study context. These differences prevent direct comparison between conditions and limit mechanistic interpretation of cross-dataset similarities or differences. Second, source localization was not performed, which restricts interpretation to scalp-level topographic dynamics rather than specific cortical or subcortical generators. Third, subjective data were limited and not temporally aligned with the EEG measures, reducing sensitivity for brain– experience associations. Fourth, only microdosing data were available for psilocybin, precluding comparison with higher-dose psilocybin effects. Finally, the naturalistic features of the DMT dataset may have introduced additional variability related to dose estimation and contextual factors.

## Conclusions

Frequency-resolved EEG microstate analysis revealed structured alterations in resting-state brain dynamics in both datasets, while broadband analysis alone captured only part of this pattern. Narrowband decomposition detected subtle low-frequency effects in the psilocybin microdosing dataset and broader multi-band alterations in the acute inhaled DMT dataset, indicating that this approach is sensitive across markedly different perturbation regimes. A descriptive overlap across datasets was observed in the delta band but should be interpreted cautiously given major design differences between the two studies. These findings support frequency-resolved microstate analysis as a useful tool for characterizing altered resting-state brain activity and for detecting frequency-specific effects not readily apparent in broadband summaries.

## Author Contributions

P.T. conceived the study, performed data analysis, and drafted the manuscript. I.G.-B. supervised the analysis, data interpretation, drafted and critically revised the manuscript. Both authors approved the final version of the manuscript.

## Competing Interests

The authors declare no competing interests.

## Acknowledgment

We would like to thank the authors of original studies, who gathered and made available the de-identified data on which this manuscript is based.

## Data Availability Statement

The EEG datasets analyzed in this study are publicly available. The psilocybin microdosing dataset can be accessed at OSF repository, and the DMT dataset is available via Zenodo. All additional data supporting the findings of this study, including processed EEG files and analysis scripts, are available from the corresponding author upon reasonable request.

## Notes

### Competing Interest Statement

The authors have declared no competing interest.

