## Supplementary Table 1 for "Characterizing Resting-State Brain Dynamics with Frequency-Resolved EEG Microstates: Parallel Analyses of Psilocybin Microdosing and Acute Inhaled DMT"

Table S1. Descriptive statistics of extracted MS parameters for each group at each frequency range for the Psilocybin dataset

| | | MS A | MS B | MS C | MS D | MS E |
|---|---|---|---|---|---|---|
| | | | | **Broadband (1-30 Hz)** | | |
| **Dur** | Psi | 49.41±3.30 | 49.40±3.93 | 67.57±13.47 | 52.76±3.61 | 51.13±3.91 |
| | Pla | 49.05±3.46 | 50.12±3.12 | 68.45±14.92 | 53.48±4.30 | 50.73±3.71 |
| **Occ** | Psi | 2.45±0.67 | 2.49±0.75 | 4.14±0.33 | 3.14±0.50 | 2.71±0.72 |
| | Pla | 2.41±0.80 | 2.53±0.69 | 3.94±0.35 | 3.31±0.57 | 2.62±0.70 |
| **Cov** | Psi | 14.01±4.77 | 14.25±5.14 | 36.18±12.26 | 19.41±4.24 | 16.15±5.65 |
| | Pla | 13.73±5.48 | 14.68±4.84 | 35.28±13.01 | 20.76±4.88 | 15.54±5.55 |
| **GEV** | Psi | 1.41±0.72 | 1.46±0.79 | 7.14±3.25 | 2.14±0.67 | 1.65±0.73 |
| | Pla | 1.63±0.85 | 1.82±0.84 | 8.36±3.74 | 2.89±1.02 | 2.04±1.16 |
| **GFP** | Psi | 1.13±0.10 | 1.13±0.11 | 1.38±0.13 | 1.18±0.11 | 1.15±0.07 |
| | Pla | 1.22±0.85 | 1.26±0.13 | 1.52±0.17 | 1.31±0.14 | 1.28±0.14 |
| | | | | **Delta (1-4 Hz)** | | |
| **Dur** | Psi | 64.72±6.05 | 65.27±5.31 | 89.96±14.02 | 66.14±3.61 | 66.59±5.08 |
| | Pla | 65.31±4.79 | 65.79±4.26 | 83.41±6.05 | 66.34±3.92 | 68.09±9.05 |
| **Occ** | Psi | 2.00±0.41 | 2.05±0.35 | 3.04±0.20 | 2.23±0.27 | 2.31±0.37 |
| | Pla | 2.10±0.35 | 2.10±0.34 | 2.97±0.26 | 2.35±0.24 | 2.40±0.33 |
| **Cov** | Psi | 15.32±4.33 | 15.81±3.64 | 33.33±6.95 | 17.34±2.71 | 18.21±4.08 |
| | Pla | 16.16±3.49 | 16.22±3.65 | 29.78±4.31 | 18.32±2.33 | 19.52±5.41 |
| **GEV** | Psi | 3.37±1.56 | 3.64±1.46 | 15.01±7.47 | 4.01±1.35 | 4.38±1.59 |
| | Pla | 4.27±1.60 | 4.45±2.05 | 13.96±3.72 | 5.21±1.68 | 5.84±2.84 |
| **GFP** | Psi | 1.13±0.11 | 1.16±0.11 | 1.45±0.20 | 1.18±0.13 | 1.19±0.12 |
| | Pla | 1.26±0.11 | 1.26±0.14 | 1.51±0.13 | 1.31±0.15 | 1.33±0.17 |
| | | | | **Theta (4-8 Hz)** | | |
| **Dur** | Psi | 47.22±2.17 | 46.25±2.14 | 61.28±7.04 | 51.93±3.02 | 48.66±2.39 |
| | Pla | 47.10±2.03 | 46.44±2.62 | 61.82±6.31 | 51.68±3.73 | 48.59±2.08 |
| **Occ** | Psi | 2.51±0.53 | 2.61±0.60 | 3.91±0.29 | 3.49±0.47 | 2.82±0.51 |
| | Pla | 2.46±0.46 | 2.56±0.54 | 3.94±0.29 | 3.49±0.54 | 2.77±0.50 |
| **Cov** | Psi | 14.08±3.43 | 14.45±3.88 | 31.97±7.02 | 22.79±4.56 | 16.71±4.09 |
| | Pla | 13.76±2.97 | 14.19±3.64 | 32.93±7.24 | 22.66±5.14 | 16.47±4.06 |
| **GEV** | Psi | 3.12±1.18 | 3.01±1.54 | 11.81±4.43 | 5.67±2.05 | 3.86±1.42 |
| | Pla | 3.83±1.49 | 3.71±1.60 | 16.12±5.89 | 7.37±2.52 | 4.89±2.08 |
| **GFP** | Psi | 1.16±0.11 | 1.13±0.13 | 1.37±0.17 | 1.23±0.11 | 1.19±0.13 |
| | Pla | 1.31±0.13 | 1.30±0.12 | 1.59±0.23 | 1.41±0.14 | 1.36±0.19 |
| | | | | **Alpha (8-13 Hz)** | | |
| **Dur** | Psi | 93.89±15.79 | 88.55±17.32 | 172.33±97.54 | 97.38±21.98 | 92.83±18.03 |
| | Ctl | 91.05±13.04 | 90.86±18.11 | 165.99±79.14 | 96.88±22.43 | 88.18±20.22 |
| **Occ** | Psi | 1.22±0.48 | 1.02±0.46 | 1.72±0.37 | 1.37±0.47 | 1.12±0.45 |
| | Ctl | 1.24±0.57 | 1.10±0.49 | 1.74±0.38 | 1.44±0.48 | 1.10±0.42 |
| **Cov** | Psi | 15.16±6.12 | 12.07±6.05 | 40.96±16.13 | 18.16±6.83 | 13.64±6.07 |
| | Ctl | 14.95±6.56 | 13.13±5.96 | 40.59±16.42 | 18.81±7.16 | 12.53±6.15 |
| **GEV** | Psi | 3.17±2.14 | 2.62±2.08 | 17.33±8.79 | 3.70±2.11 | 2.80±1.72 |
| | Ctl | 3.39±1.91 | 3.25±2.13 | 20.80±10.39 | 4.79±3.19 | 3.23±2.77 |
| **GFP** | Psi | 1.21±0.27 | 1.20±0.29 | 1.57±0.34 | 1.23±0.26 | 1.22±0.26 |
| | Ctl | 1.29±0.21 | 1.32±0.29 | 1.72±0.38 | 1.34±0.28 | 1.30±0.25 |
| | | | | **Beta (13-30 Hz)** | | |
| **Dur** | Psi | 64.46±5.50 | 68.64±4.16 | 81.77±11.26 | 67.83±7.05 | 61.96±5.18 |
| | Ctl | 64.46±4.03 | 68.80±3.77 | 80.79±9.80 | 68.22±6.31 | 61.86±4.88 |
| **Occ** | Psi | 2.14±0.42 | 2.58±0.44 | 3.28±0.36 | 2.36±0.42 | 1.84±0.41 |
| | Ctl | 2.16±0.46 | 2.62±0.36 | 3.23±0.28 | 2.40±0.40 | 1.81±0.40 |
| **Cov** | Psi | 15.79±3.75 | 20.52±3.74 | 32.06±5.57 | 18.66±4.63 | 12.96±3.54 |
| | Ctl | 15.96±4.21 | 20.83±3.01 | 31.32±5.92 | 19.11±4.61 | 12.78±3.44 |
| **GEV** | Psi | 3.20±1.13 | 4.47±1.38 | 11.26±4.01 | 3.77±1.42 | 2.79±0.97 |
| | Ctl | 3.27±1.14 | 4.70±0.95 | 11.28±3.24 | 3.93±1.39 | 2.92±1.31 |
| **GFP** | Psi | 1.20±0.13 | 1.22±0.12 | 1.37±0.15 | 1.23±0.13 | 1.21±0.14 |
| | Ctl | 1.20±0.10 | 1.23±0.09 | 1.37±0.12 | 1.22±0.11 | 1.21±0.13 |
