## Supplementary Table 2 for "Characterizing Resting-State Brain Dynamics with Frequency-Resolved EEG Microstates: Parallel Analyses of Psilocybin Microdosing and Acute Inhaled DMT"

Table S2. Descriptive statistics of extracted MS parameters for each group at each frequency range for the DMT dataset

| | | MS A | MS B | MS C | MS D | MS E | MS F |
|---|---|---|---|---|---|---|---|
| **Broadband (1-30 Hz)** | | | | | | | |
| **Dur** | Post | 50.79±4.30 | 50.15±5.78 | 62.80±7.03 | 53.30±3.26 | 52.01±4.67 | N.A. |
| | Pre | 50.25±3.29 | 50.42±2.92 | 64.97±9.25 | 54.57±3.19 | 52.06±3.96 | N.A. |
| **Occ** | Post | 2.71±0.74 | 2.55±0.96 | 4.15±0.74 | 3.22±0.65 | 2.77±0.78 | N.A. |
| | Pre | 2.50±0.49 | 2.51±0.63 | 4.06±0.45 | 3.35±0.60 | 2.77±0.67 | N.A. |
| **Cov** | Post | 15.97±5.96 | 15.32±8.00 | 31.88±9.32 | 19.90±5.62 | 16.93±6.53 | N.A. |
| | Pre | 14.33±3.23 | 14.52±4.44 | 32.84±8.67 | 21.45±5.00 | 16.86±5.21 | N.A. |
| **GEV** | Post | 2.56±1.54 | 2.49±1.85 | 9.52±4.83 | 3.62±2.33 | 2.99±2.06 | N.A. |
| | Pre | 3.32±1.17 | 3.61±154 | 15.64±8.40 | 6.03±2.44 | 4.34±2.02 | N.A. |
| **GFP** | Post | 1.18±0.17 | 1.18±0.16 | 1.39±0.26 | 1.22±0.25 | 1.18±0.16 | N.A. |
| | Pre | 1.30±0.22 | 1.33±0.23 | 1.56±0.32 | 1.39±0.25 | 1.35±0.23 | N.A. |
| **Delta (1-4 Hz)** | | | | | | | |
| **Dur** | Post | 64.08±10.26 | 69.07±13.80 | 96.19±16.66 | 68.37±6.09 | 69.33±7.05 | N.A. |
| | Pre | 63.37±5.20 | 66.01±5.54 | 85.94±7.03 | 67.46±3.96 | 68.06±5.29 | N.A. |
| **Occ** | Post | 1.62±0.58 | 1.85±0.68 | 3.04±0.52 | 2.10±0.40 | 2.06±0.46 | N.A. |
| | Pre | 1.94±0.26 | 2.15±0.27 | 3.05±0.26 | 2.35±0.26 | 2.30±0.38 | N.A. |
| **Cov** | Post | 12.85±6.82 | 16.19±9.58 | 36.46±11.04 | 17.17±4.71 | 17.33±5.49 | N.A. |
| | Pre | 14.36±3.00 | 16.67±3.40 | 31.70±5.09 | 18.66±2.87 | 18.60±4.23 | N.A. |
| **GEV** | Post | 3.19±2.87 | 4.56±4.43 | 17.12±8.76 | 4.08±2.30 | 4.26±2.70 | N.A. |
| | Pre | 2.84±1.17 | 3.39±1.50 | 11.41±5.50 | 3.78±1.29 | 3.90±1.54 | N.A. |
| **GFP** | Post | 1.38±0.27 | 1.42±0.27 | 1.76±0.47 | 1.44±0.48 | 1.40±0.24 | N.A. |
| | Pre | 1.15±0.11 | 1.16±0.12 | 1.39±0.15 | 1.18±0.09 | 1.19±0.09 | N.A. |
| **Theta (4-8 Hz)** | | | | | | | |
| **Dur** | Post | 47.66±3.65 | 45.36±3.63 | 57.36±5.13 | 50.59±2.73 | 47.19±3.53 | N.A. |
| | Pre | 48.06±2.21 | 46.69±2.10 | 58.40±6.35 | 52.89±3.27 | 48.24±2.83 | N.A. |
| **Occ** | Post | 2.86±0.61 | 2.50±0.81 | 4.00±0.62 | 3.59±0.64 | 2.98±0.71 | N.A. |
| | Pre | 2.82±0.51 | 2.62±0.53 | 3.74±0.29 | 3.55±0.44 | 2.89±0.73 | N.A. |
| **Cov** | Post | 16.57±5.57 | 13.71±5.80 | 29.69±7.45 | 22.47±5.35 | 17.55±5.97 | N.A. |
| | Pre | 16.29±3.42 | 14.55±3.40 | 28.49±6.30 | 23.63±4.34 | 17.05±5.28 | N.A. |
| **GEV** | Post | 3.63±2.22 | 2.58±1.76 | 10.28±5.28 | 5.20±2.72 | 3.78±2.99 | N.A. |
| | Pre | 4.95±1.89 | 3.94±1.83 | 13.38±6.81 | 8.06±3.73 | 4.79±2.16 | N.A. |
| **GFP** | Post | 1.22±0.16 | 1.18±0.15 | 1.41±0.22 | 1.26±0.20 | 1.21±0.18 | N.A. |
| | Pre | 1.38±0.31 | 1.33±0.31 | 1.57±0.42 | 1.45±0.38 | 1.36±0.32 | N.A. |
| **Alpha (8-13 Hz)** | | | | | | | |
| **Dur** | Post | 86.10±13.19 | 76.11±9.33 | 97.91±12.75 | 82.89±12.27 | 77.99±14.23 | 70.03±10.06 |
| | Pre | 96.25±16.24 | 91.11±16.78 | 130.74±38.24 | 102.76±19.03 | 93.56±20.17 | 75.77±11.04 |
| **Occ** | Post | 1.64±0.35 | 1.30±0.28 | 1.86±0.39 | 1.79±0.34 | 1.46±0.41 | 1.21±0.31 |
| | Pre | 1.21±0.36 | 1.04±0.29 | 1.64±0.28 | 1.51±0.40 | 1.18±0.35 | 0.84±0.33 |
| **Cov** | Post | 18.22±6.08 | 12.65±3.91 | 23.86±6.79 | 19.66±5.49 | 14.74±5.68 | 10.87±4.17 |
| | Pre | 15.01±5.04 | 12.35±4.50 | 28.87±9.52 | 21.07±6.92 | 14.62±5.40 | 8.08±3.34 |
| **GEV** | Post | 2.60±1.85 | 1.50±0.95 | 5.18±3.30 | 2.48±.1.61 | 1.68±1.40 | 1.01±0.85 |
| | Pre | 6.32±3.04 | 4.93±2.72 | 20.27±9.90 | 8.22±4.32 | 4.96±2.90 | 1.95±1.13 |
| **GFP** | Post | 1.20±0.09 | 1.15±0.12 | 1.39±0.20 | 1.19±0.16 | 1.12±0.12 | 1.07±0.10 |
| | Pre | 2.09±0.62 | 2.06±0.59 | 2.52±0.82 | 2.11±0.65 | 1.99±0.63 | 1.77±0.47 |
| **Beta (1-4 Hz)** | | | | | | | |
| **Dur** | Post | 59.71±4.22 | 59.45±3.32 | 65.69±7.61 | 61.87±5.67 | 60.08±8.33 | 57.40±5.26 |
| | Pre | 61.13±4.07 | 62.08±4.40 | 68.03±7.16 | 66.66±6.84 | 61.36±5.69 | 57.42±3.97 |
| **Occ** | Post | 2.37±0.57 | 2.29±0.46 | 2.70±0.64 | 2.58±0.59 | 2.02±0.68 | 2.10±0.75 |
| | Pre | 2.12±0.32 | 2.28±0.45 | 2.73±0.45 | 2.68±0.44 | 2.12±0.53 | 1.79±0.46 |
| **Cov** | Post | 16.14±4.78 | 15.46±3.79 | 21.04±8.07 | 18.51±5.93 | 14.87±7.36 | 13.99±7.06 |
| | Pre | 14.66±2.86 | 16.14±4.01 | 21.65±5.24 | 20.91±5.05 | 15.08±5.31 | 11.57±3.47 |
| **GEV** | Post | 3.39±1.29 | 3.54±1.82 | 6.24±3.25 | 3.50±1.67 | 2.77±2.45 | 2.56±2.15 |
| | Pre | 3.64±1.50 | 4.24±1.69 | 8.01±3.80 | 4.92±2.15 | 3.14±1.52 | 2.10±0.99 |
| **GFP** | Post | 1.29±0.20 | 1.31±0.26 | 1.38±0.22 | 1.29±0.23 | 1.23±0.21 | 1.27±0.26 |
| | Pre | 1.27±0.15 | 1.31±0.17 | 1.40±0.19 | 1.30±0.17 | 1.26±0.18 | 1.22±0.15 |
