## Supplementary Table 3 for "Characterizing Resting-State Brain Dynamics with Frequency-Resolved EEG Microstates: Parallel Analyses of Psilocybin Microdosing and Acute Inhaled DMT"

*Table S3. Correlation coefficients and corresponding p values between MS parameters and VAS scores in the Psilocybin condition of the Psilocybin dataset*

| | **GEV** | | | | | **GFP** | | | | | **Duration** | | | | | **Coverage** | | | | | **Occurrence** | | | | |
|---|---|---|---|---|---|---|---|---|---|---|---|---|---|---|---|---|---|---|---|---|---|---|---|---|---|
| | *MS A* | *MS B* | *MS C* | *MS D* | *MD E* | *MS A* | *MS B* | *MS C* | *MS D* | *MD E* | *MS A* | *MS B* | *MS C* | *MS D* | *MD E* | *MS A* | *MS B* | *MS C* | *MS D* | *MD E* | *MS A* | *MS B* | *MS C* | *MS D* | *MD E* |
| **Broadband (1-30 Hz)** | | | | | | | | | | | | | | | | | | | | | | | | | |
| *p* | 0.909 | 0.373 | 0.496 | 0.318 | 0.382 | 0.318 | 0.318 | 0.582 | 0.318 | 0.897 | 0.318 | 0.368 | 0.401 | 0.318 | 0.382 | 0.318 | 0.382 | 0.382 | 0.318 | 0.401 | 0.318 | 0.401 | 0.318 | 0.401 | 0.409 |
| *r* | -0.021 | 0.248 | -0.145 | 0.44 | -0.218 | 0.313 | 0.294 | 0.116 | 0.356 | -0.033 | 0.312 | 0.258 | -0.195 | 0.292 | -0.215 | 0.307 | 0.226 | -0.228 | 0.293 | -0.19 | 0.284 | 0.19 | -0.28 | 0.184 | -0.176 |
| **Delta (1-4 Hz)** | | | | | | | | | | | | | | | | | | | | | | | | | |
| *p* | 0.986 | 0.98 | 0.98 | 0.35 | 0.98 | 0.986 | 0.98 | 0.98 | 0.975 | 0.986 | 0.986 | 0.986 | 0.986 | 0.986 | 0.35 | 0.986 | 0.986 | 0.986 | 0.659 | 0.975 | 0.986 | 0.986 | 0.975 | 0.659 | 0.98 |
| *r* | 0.077 | 0.123 | 0.152 | 0.395 | -0.145 | 0.107 | 0.128 | 0.162 | 0.22 | -0.026 | -0.044 | -0.067 | 0.048 | 0.082 | -0.395 | 0.003 | 0.024 | 0.005 | 0.296 | -0.23 | 0.005 | 0.04 | -0.203 | 0.302 | -0.13 |
| **Theta (4-8 Hz)** | | | | | | | | | | | | | | | | | | | | | | | | | |
| *p* | 0.636 | 0.791 | 0.636 | 0.636 | 0.662 | 0.737 | 0.865 | 0.642 | 0.938 | 0.636 | 0.865 | 0.79 | 0.636 | 0.636 | 0.737 | 0.662 | 0.79 | 0.636 | 0.636 | 0.737 | 0.662 | 0.737 | 0.636 | 0.636 | 0.662 |
| *r* | 0.246 | 0.073 | -0.254 | 0.215 | -0.158 | 0.12 | -0.048 | -0.199 | -0.015 | -0.223 | 0.04 | -0.081 | -0.265 | 0.265 | -0.11 | 0.179 | 0.081 | -0.262 | 0.32 | -0.134 | 0.175 | 0.109 | -0.211 | 0.249 | -0.158 |
| **Alpha (8-13 Hz)** | | | | | | | | | | | | | | | | | | | | | | | | | |
| *p* | 0.567 | 0.627 | 0.842 | 0.567 | 0.627 | 0.176 | 0.176 | 0.176 | 0.176 | 0.567 | 0.714 | 0.714 | 0.627 | 0.906 | 0.631 | 0.567 | 0.567 | 0.567 | 0.567 | 0.631 | 0.627 | 0.627 | 0.925 | 0.567 | 0.739 |
| *r* | 0.272 | 0.175 | -0.054 | 0.241 | -0.149 | 0.418 | 0.404 | 0.394 | 0.449 | 0.239 | 0.099 | 0.098 | -0.157 | -0.031 | -0.139 | 0.231 | 0.206 | -0.211 | 0.226 | -0.132 | 0.148 | 0.16 | 0.018 | 0.203 | -0.085 |
| **Beta (13-30 Hz)** | | | | | | | | | | | | | | | | | | | | | | | | | |
| *p* | 0.397 | 0.493 | 0.74 | 0.953 | 0.784 | 0.493 | 0.493 | 0.603 | 0.603 | 0.603 | 0.397 | 0.493 | 0.884 | 0.784 | 0.884 | 0.397 | 0.603 | 0.578 | 0.784 | 0.689 | 0.493 | 0.74 | 0.493 | 0.689 | 0.603 |
| *r* | 0.372 | 0.267 | -0.113 | 0.011 | -0.089 | 0.27 | 0.317 | 0.182 | 0.203 | 0.17 | 0.359 | 0.295 | -0.036 | 0.075 | 0.039 | 0.369 | 0.186 | -0.222 | -0.08 | -0.135 | 0.249 | 0.108 | -0.258 | -0.143 | -0.169 |
