## Supplementary Table 4 for "Characterizing Resting-State Brain Dynamics with Frequency-Resolved EEG Microstates: Parallel Analyses of Psilocybin Microdosing and Acute Inhaled DMT"

*Table S4. Correlation coefficients and corresponding p values between MS parameters and 5D-ASC scores in the DMT dataset*

| | GEV | | | | | | GFP | | | | | | Duration | | | | | | Coverage | | | | | | Occurrence | | | | | |
|---|---|---|---|---|---|---|---|---|---|---|---|---|---|---|---|---|---|---|---|---|---|---|---|---|---|---|---|---|---|---|
| | *MS A* | *MS B* | *MS C* | *MS D* | *MD E* | *MS F* | *MS A* | *MS B* | *MS C* | *MS D* | *MD E* | *MS F* | *MS A* | *MS B* | *MS C* | *MS D* | *MD E* | *MS F* | *MS A* | *MS B* | *MS C* | *MS D* | *MD E* | *MS F* | *MS A* | *MS B* | *MS C* | *MS D* | *MD E* | *MS F* |
| **Broadband (1-30 Hz)** | | | | | | | | | | | | | | | | | | | | | | | | | | | | | | |
| *p* | 0.99 | 0.99 | 0.99 | 0.99 | 0.99 | N.A. | 0.99 | 0.99 | 0.99 | 0.99 | 0.99 | N.A. | 0.99 | 0.99 | 0.99 | 0.99 | 0.99 | N.A. | 0.99 | 0.99 | 0.99 | 0.99 | 0.99 | N.A. | 0.99 | 0.99 | 0.99 | 0.99 | 0.99 | N.A. |
| *r* | 0.169 | 0.137 | 0.174 | 0.068 | -0.002 | N.A. | 0.223 | 0.05 | 0.209 | 0.104 | 0.06 | N.A. | -0.008 | 0.122 | -0.035 | -0.125 | -0.23 | N.A. | 0.026 | -0.017 | 0.09 | -0.084 | -0.149 | N.A. | 0.159 | 0.055 | 0.274 | 0.061 | -0.04 | N.A. |
| **Delta (1-4 Hz)** | | | | | | | | | | | | | | | | | | | | | | | | | | | | | | |
| *p* | 0.783 | 0.783 | 0.783 | 0.783 | 0.783 | N.A. | 0.783 | 0.783 | 0.783 | 0.783 | 0.783 | N.A. | 0.783 | 0.952 | 0.783 | 0.783 | 0.783 | N.A. | 0.795 | 0.936 | 0.783 | 0.783 | 0.783 | N.A. | 0.783 | 0.783 | 0.783 | 0.783 | 0.783 | N.A. |
| *r* | 0.202 | 0.227 | 0.182 | 0.144 | 0.105 | N.A. | 0.082 | 0.17 | 0.099 | 0.078 | 0.096 | N.A. | -0.119 | 0.011 | -0.216 | -0.182 | -0.114 | N.A. | -0.061 | 0.023 | -0.085 | -0.087 | -0.071 | N.A. | 0.158 | 0.195 | 0.208 | 0.126 | 0.078 | N.A. |
| **Theta (4-8 Hz)** | | | | | | | | | | | | | | | | | | | | | | | | | | | | | | |
| *p* | 0.944 | 0.799 | 0.969 | 0.944 | 0.944 | N.A. | 0.799 | 0.799 | 0.969 | 0.944 | 0.944 | N.A. | 0.944 | 0.944 | 0.944 | 0.944 | 0.944 | N.A. | 0.799 | 0.799 | 0.944 | 0.944 | 0.969 | N.A. | 0.944 | 0.799 | 0.944 | 0.944 | 0.961 | N.A. |
| *r* | 0.175 | 0.302 | 0.008 | 0.047 | 0.083 | N.A. | 0.255 | 0.355 | 0.007 | 0.059 | 0.076 | N.A. | 0.115 | 0.171 | -0.055 | -0.072 | -0.116 | N.A. | 0.26 | 0.229 | 0.07 | 0.05 | 0.008 | N.A. | 0.073 | 0.241 | -0.097 | -0.055 | -0.035 | N.A. |
| **Alpha (8-13 Hz)** | | | | | | | | | | | | | | | | | | | | | | | | | | | | | | |
| *p* | 0.997 | 0.997 | 0.997 | 0.997 | 0.997 | 0.997 | 0.997 | 0.997 | 0.997 | 0.997 | 0.997 | 0.997 | 0.997 | 0.997 | 0.997 | 0.997 | 0.997 | 0.997 | 0.997 | 0.997 | 0.997 | 0.997 | 0.997 | 0.997 | 0.997 | 0.997 | 0.997 | 0.997 | 0.997 | 0.997 |
| *r* | -0.018 | 0.043 | -0.163 | -0.158 | -0.044 | 0.145 | 0.191 | 0.008 | -0.001 | 0.068 | 0.242 | 0.303 | -0.024 | 0.038 | -0.036 | 0.234 | 0.107 | 0.017 | 0.01 | -0.025 | 0.176 | 0.236 | 0.09 | 0.048 | -0.003 | -0.018 | 0.043 | -0.163 | -0.158 | -0.044 |
| **Beta (13-30 Hz)** | | | | | | | | | | | | | | | | | | | | | | | | | | | | | | |
| *p* | 0.92 | 0.907 | 0.92 | 0.907 | 0.907 | 0.907 | 0.907 | 0.92 | 0.92 | 0.945 | 0.945 | 0.907 | 0.92 | 0.907 | 0.907 | 0.907 | 0.907 | 0.92 | 0.92 | 0.945 | 0.907 | 0.907 | 0.907 | 0.92 | 0.92 | 0.92 | 0.907 | 0.92 | 0.907 | 0.907 |
| *r* | 0.039 | 0.245 | -0.066 | -0.116 | -0.137 | 0.135 | 0.194 | 0.086 | 0.039 | 0.021 | 0.012 | 0.24 | -0.044 | -0.121 | -0.211 | 0.165 | 0.212 | 0.069 | 0.042 | 0.017 | 0.161 | 0.24 | 0.154 | 0.04 | -0.091 | 0.039 | 0.245 | -0.066 | -0.116 | -0.137 |
